# *TreeCompR*: Tree competition indices for inventory data and 3D point clouds

**DOI:** 10.1101/2024.03.23.586379

**Authors:** Julia S. Rieder, Roman M. Link, Konstantin Köthe, Dominik Seidel, Tobias Ullmann, Anja Žmegač, Christian Zang, Bernhard Schuldt

**Author notes:** First and second author contributed equally.

## Abstract

1. In times of more frequent global-change-type droughts and associated tree mortality events, competition release is one silvicultural measure discussed to have an impact on the resilience of managed forest stands. Understanding how trees compete with each other is therefore crucial, but different measurement options and competition indices leave users with the agony of choice, as no single competition index has proven universally superior.
2. To help users with the choice and computation of appropriate indices, we present the open-source *TreeCompR* package, which can handle 3D point clouds in various formats as well as classical forest inventory data and serves as a centralized platform for exploring and comparing different competition indices (CIs). Within a common interface, users can efficiently select the most suitable CI for their specific research questions. The package facilitates the integration of both traditional distance-dependent and novel point cloud-based indices.
3. To evaluate the package, we used *TreeCompR* to quantify the competition situation of 308 European beech trees from 13 sites in Central Europe. Based on this dataset, we discuss the interpretation, comparability and sensitivity of the different indices to their parameterization and identify possible sources of uncertainty and ways to minimize them.
4. The compatibility of *TreeCompR* with different data formats and different data collection methods makes it accessible and useful for a wide range of users, specifically ecologists and foresters. Due to the flexibility in the choice of input formats as well as the emphasis on tidy, well-structured output, our package can easily be integrated into existing data-analysis workflows both for 3D point cloud and classical forest inventory data.

## 1. Introduction

In the wake of climate-change driven mass tree mortality events in Europe (Schuldt et al., 2020) and elsewhere (Hammond et al., 2022), there is a growing interest in the biotic and abiotic factors that influence tree vitality (Cailleret et al., 2017; Chakraborty et al., 2017). This is specifically true for competition, which is directly amenable to silvicultural measures such as thinning and therefore important for forest management (cf. Bose et al. 2021). For these reasons, there is an urgent need for rapid, reproducible and easy-to-use tools to quantify tree competition.

Competition indices (CI) are commonly used metrics that allow to quantify tree competition and resource availability for individual trees within a stand. CIs provide a single value per tree that serves as a general indicator to which extent a tree has to share water, light, nutrients and particularly its growing space with neighbouring trees (Bartsch & Röhrig, 2016). The nature of competition can also vary depending on the location. In areas with limited resources, light may not be a primary constraint, as tree growth likewise is impeded by water and nutrient availability (Pretzsch, 2019).

Numerous traditional methods exist for quantifying tree competition. Weigelt & Jolliffe (2003) and Contreras et al. (2011) compared these different CIs and concluded that no universally applicable competition index exists. Traditional approaches are based on the measurement of classical forest inventory data such as the diameter at breast height (DBH), tree height (H) and the position of the target trees and their neighbours in the field. Such inventory data can be used to calculate, for example, the Hegyi index (1974) or other size-ratio indices (Contreras et al., 2011).

Although field inventories provide detailed information, they are time consuming and subject to observer bias. To overcome these challenges, airborne and ground-based light detection and ranging (LiDAR) technologies have proven to be a valuable tool for forest analysis. They provide 3D point clouds of objects from different platforms and allow a flexible choice of method depending on research needs and desired point density, with point density/resolution increasing from space-based to ground-based detection (Dassot et al., 2011; Calders et al., 2020).

The structural parameters that are traditionally collected in the field as inventory data can also be derived from LiDAR-based point clouds. R packages that focus on the segmentation and derivation of structural parameters include, e.g., *lidR* (Roussel et al., 2020) for airborne data, *TreeLS* (de Conto et al., 2017) and *FORTLS* (Molina-Valero et al., 2022) for terrestrial laser scanning (TLS) or mobile laser scanning (MLS) data or *ITSMe* (Terryn et al., 2022) for individual tree point clouds. Additionally, the R package *forestecology* is available for tree growth models (Kim et al., 2021). While all these packages help obtaining the data necessary to calculate CIs, no package is available so far that automates their computation.

Typically, in competition modelling where crown metrics are considered, geometric standard crown shapes are employed (Pretzsch et al., 2002), while conventional methods for evaluating crown parameters are both labour-intensive and prone to measurement mistakes and thus are rarely included in crown indices (Bayer et al., 2013; von Oheimb et al., 2011). Laser scanning (TLS/MLS) can be less affected by observer effects and was shown to be as reliable as traditional methods, e.g., compared to eight-point crown projections (Calders et al., 2020; Seidel et al., 2015).

Addressing the challenge of varying tree shapes within the same species and topographic effects at plot level, recent studies proposed an approach directly working with the LiDAR point clouds for quantifying competition (Metz et al., 2013; Seidel et al., 2015). These studies used an adapted version of the KKL index (‘Kronenkonkurrenz um Licht’, canopy competition for light) by Pretzsch (1995) and Pretzsch et al. (2002), spanning a virtual search cone at the target tree’s position, determining how much of the points or voxels belonging to neighbouring tree crowns extend into this cone. The higher this count, the stronger the competition, with a freely standing tree having a count of 0 within the cone. This approach can also be applied with a search cylinder (Seidel et al., 2015) around the target tree instead of a cone, to also include competition by lower trees or understory.

The high number of available competition metrics and data collection options complicates selecting the most useful index for a particular research question. To fill this gap, we introduce the R package *TreeCompR*, a useful tool to overcome the challenges of navigating through different indices and their uncertainties. Our package does not only offer an overview of commonly used competition indices, but also allows users to choose a method depending on data availability or targeted research questions (target variable, respectively) and provides fast and direct outputs for specific competition measures. In the following, we will demonstrate the package’s functionality through application examples and a comparison of point cloud-based and distance-dependent indices. Additionally, we evaluate possible uncertainties and ways to overcome them.

## 2. Materials and Methods

### 2.1 Package availability and overview

*TreeCompR* has been developed to automatically calculate different competition indices based on inventory or laser scanning data. The current version (0.0.0.9000) of the *TreeCompR* package is freely available on Github (https://github.com/juliarieder/TreeCompR). The package requires R version >= 3.5.0. Detailed documentation and usage examples of all functions are provided in the package manual, and additional information is available in the vignettes and on Github.

An overview of the available methods within the *TreeCompR* package depending on the input data type is given in Figure 1. The left part of Figure 1 shows the two methods that can be used within the function compete_pc(), which calculates CIs directly from the original 3D point clouds, where only the target tree needs to be (manually) segmented beforehand. Point clouds can be read from various data formats including.las/.laz as well as a large number of possible plain text formats. The right part shows which functions can be used with inventory data from all kinds of collection sources, and which steps would be needed to derive the inventory table from point clouds, depending on the collection type. The function compete_inv() can compute three different distance-DBH-dependent as well as three distance-height-dependent CIs. If method is set to "all_methods", all CIs will be derived that can be calculated from the provided input. For example, for airborne laser scanning (ALS) data (where due to the nature of the data no DBH values are available) only distance-height-dependent CIs will be calculated. ALS/TLS or MLS data can be used as a source of inventory data but require pre-processing through a full (automated) segmentation of individual trees within a plot or the extraction of tree heights from a canopy height model (CHM). The functions read_pc() and read_inv() provide flexible parsers for both point cloud- and inventory-based datasets in a variety of formats that are used internally in compete_pc() and read_pc(). Table 1 shows in more detail which CIs can be quantified with the *TreeCompR* package, how each index is calculated, which *TreeCompR* functions can be used to compute them, and which are the most relevant references for each index.

**Figure 1:**
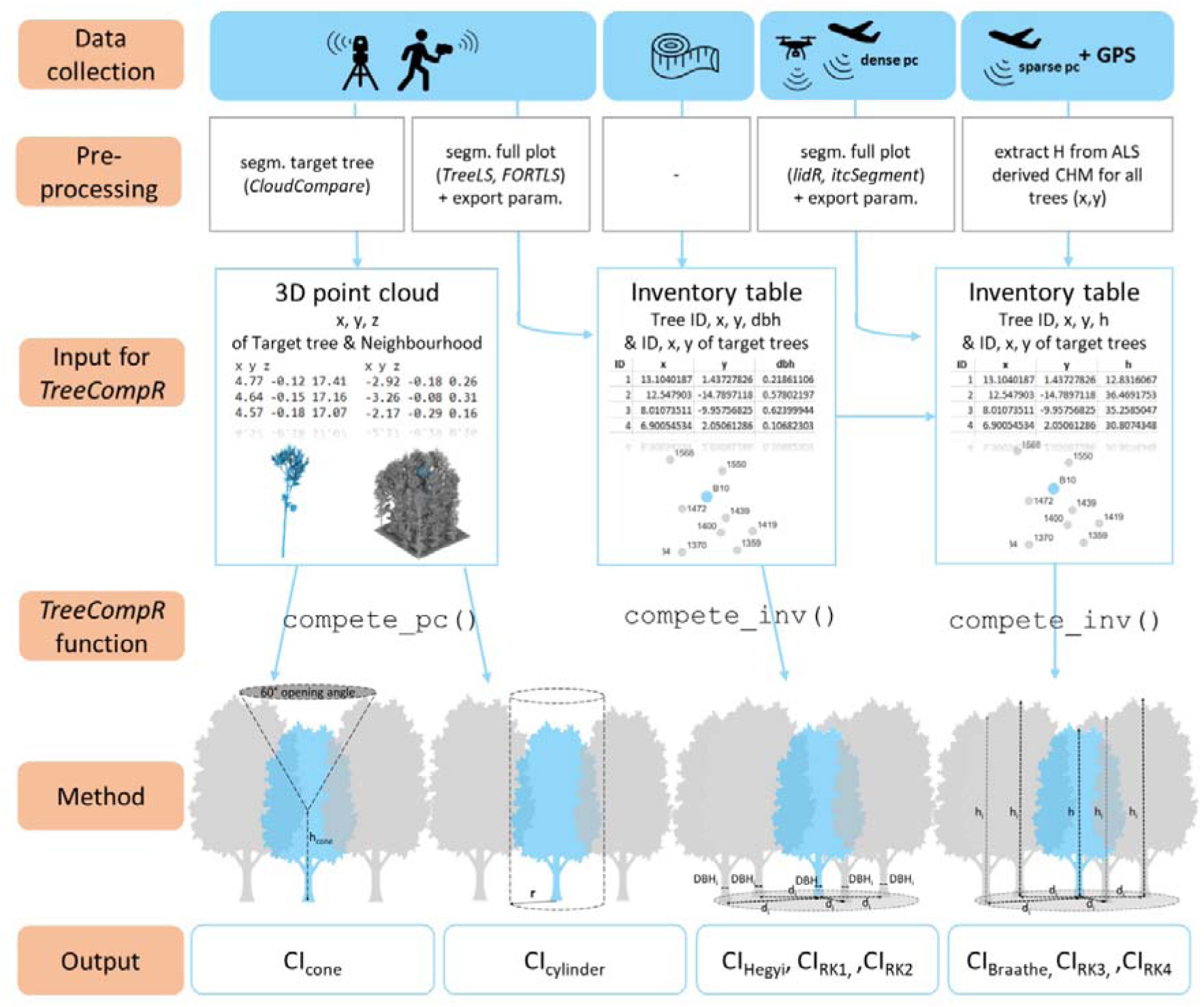
Methodological overview within *TreeCompR* – left: functions and competition indices (CIs) related to input of original point clouds from close-range remote sensing as terrestrial or mobile laser scanning (TLS/MLS); right: functions and CIs related to input of inventory data that can be derived by various data collection methods as ground-based inventory, TLS/MLS or airborne laser scanning (ALS). R package recommendations for pre-processing the point clouds are written in italics.

**Table 1:** List of the competition indices (CIs) available in the *TreeCompR* package, the associated formula, the corresponding function name and literature reference, with diameter at breast height (DBH, m) and height (h, m) of the target tree, diameter at breast height (DBH_i_, m) and height (h_i_, m) of competitor i, and the distance between target tree and competitor i (d_i_, m)

| CI | Description/Equation | Function in *TreeCompR* | Reference |
| --- | --- | --- | --- |
| $CI_{\text{cone}}$ | $n_{\text{voxel}}$ of competing neighbourhood within cone | `compete_pc()` | Pretzsch (1995), Pretzsch et al. (2002), Metz et al. (2013), Seidel et al. (2015) |
| $CI_{\text{cylinder}}$ | $n_{\text{voxel}}$ of competing neighbourhood within cylinder | `compete_pc()` | Seidel et al. (2015) |
| $CI_{\text{Hegyi}}$ | $\sum_{i=1}^n DBH_i / (DBH \cdot d_i)$ | `compete_inv()` | Hegyi (1974) |
| $CI_{\text{Braathe}}$ | $\sum_{i=1}^n h_i / (h \cdot d_i)$ | `compete_inv()` | Braathe (1980) |
| $CI_{\text{RK1}}$ | $\sum_{i=1}^n \arctan(DBH_i / d_i)$ | `compete_inv()` | Rouvinen and Kuuluvainen (1997) |
| $CI_{\text{RK2}}$ | $\sum_{i=1}^n \left( \frac{DBH_i}{DBH} \right) \cdot \arctan(DBH_i / d_i)$ | `compete_inv()` | Rouvinen and Kuuluvainen (1997) |
| $CI_{\text{RK3}}$ | $\sum_{i=1}^n \arctan(h_i / d_i), h_i > h$ | `compete_inv()` | Rouvinen and Kuuluvainen (1997) |
| $CI_{\text{RK4}}$ | $\sum_{i=1}^n \left( \frac{h_i}{h} \right) \cdot \arctan(h_i / d_i)$ | `compete_inv()` | Rouvinen and Kuuluvainen (1997) |

### 2.2 Field data collection and pre-processing

We use example data derived from hand-held mobile laser scans (MLS) conducted in 13 forest sites distributed over northern Bavaria, Germany, in Central Europe to demonstrate the functionality of the *TreeCompR* package. The scans were carried out using a GeoSLAM ZEB Horizon (GeoSLAM Ltd., Nottingham, UK) in the leaf-off period in March/April 2022 to obtain the 3D tree structures. As target trees, the dataset encompasses 308 mature European beech trees (*Fagus sylvatica* L.) of different crown vitality levels. All 13 sites were affected by the extreme drought in 2018/19 in Central Europe (Schuldt et al., 2020), and show differences in tree vitality (crown defoliation) within each site. The inclusion of pure and mixed forests, as well as managed and unmanaged areas, ensures a broad representation of ecological conditions.

Crown projection area (CPA, m^2^) and diameter at breast height (DBH, m) of each target tree were determined using the *ITSMe* package v1.0.0 (Terryn et al., 2022), and tree height (H, m) was calculated with tree_pos()from our package. Additionally, the box-dimension (D_b_) was calculated using the *VoxR* package v1.0.0 (Lecigne et al., 2018). D_b_ is an integrative measure of the structural complexity of a tree (Seidel, 2018; Saarinen et al., 2021). Its values range between 1 (pole-like) and 3 (solid cube), though there is an upper limit somewhere below 2.72 for trees (Seidel et al., 2019). These variables are associated with the vitality and structure of trees and will be examined to assess how well the competition indices capture differences in ecological attributes.

Prior to quantifying distance-dependent CIs, we used the *TreeLS* R package v2.0.5 (de Conto et al., 2017) to segment the plots from MLS point clouds and extract individual tree information. The inventory table needed to compute these indices with compete_inv()needs to contain cartesian spatial coordinates (x, y) in metres, and, depending on the index, either DBH (in cm) or height (in m). If different units are used, this can be specified within the function parameters in compete_inv().

In addition to the MLS data, we tested the package with publicly available ALS data for all forest sites. The ALS data were obtained by the Bavarian agency for digitisation, high-speed internet and surveying (Landesamt für Digitalisierung, Breitband und Vermessung, LDBV) in 2022 for the years 2010-2014, containing first and last returns (min. point density 4 pts/m^2^). The ALS point clouds were segmented using R packages *lidR* v4.0.4 and *itcSegment* v1.0 (cf. Figure 1).

### 2.3 Competition indices based on inventory data

With the function compete_inv(), we quantified the distance-dependent indices as described in Table 1. The CI_Hegyi_ is the weighted sum of the relative diameters of the neighbours in the search radius (relative to the diameter of the target tree) weighted by the inverse distance to the target tree (Table 1). The CI_Braathe_ is a modification of this index based on relative height instead of relative diameter (Table 1). The CI_RK1_ to CI_RK4_ according to Rouvinen and Kuuluvainen (1997) include the sums of the subtended angles and include DBH or height, each weighted by inverse distance and, in case of CI_RK2_ and CI_RK4_, their relative size. In addition, CI_RK3_ only includes neighbouring trees as competitors if they are taller than the target tree. Users can choose whether to quantify CIs only for the trees that are not situated on the plot margins (target_source = "buff_edge" or "exclude_edge"), all trees within a plot (target source = "all_trees"), or, as in our example, for a manually specified list of target trees. To check which trees are included as target trees, users can plot a map of the defined target trees, the search radius and the included competitors with plot_target(). If no target trees are specified, by default the trees closer than one search radius to the edge of the plot (approximated by a concave hull) are excluded from the list of target trees, as competition for these positions is underestimated due to edge effects. If the plot edge is a natural forest edge, it is also possible to run the analysis for all trees.

Within compete_inv(), the user can adjust the search radius (in m) around the target trees where neighbours are considered as competitors. This setting is particularly important as the variability in crown size between different tree species or age classes should be taken into account. Lorimer (1983) suggests that the search radius for competitors should be set at 3.5-times the mean crown radius of the target trees. In our example, we set the search radius to 13.5 m, which is three times the crown radius (4.5 m). Higher values were not possible because the point density was lower at the edges of our plots and we wanted to ensure consistent data quality across our datasets. As all distance-based indices are very sensitive to this radius, a deliberate and informed decision for an appropriate search radius prior to the analysis is of utmost importance to ensure replicable and meaningful results.

### 2.4 Competition indices based on 3D point cloud data

For quantifying the competition from point clouds, the target trees were manually segmented in *CloudCompare* (v2.11.3, https://www.danielgm.net/cc/) to assure branches are not wrongly assigned to other trees. The point clouds of segmented target trees and 30×30 m plot files centred around the target tree were exported from *CloudCompare*. The functionality of compete_pc() is illustrated in Figure 2. In both method cases, the point cloud of the target tree and of its plot need to be specified. Within the function, the competing neighbourhood is filtered from the plot. As laser scanning from the ground can lead to occlusion effects in the upper crown (cf. Mathes et al., 2023), some crown parts might have a lower point density in the datasets. To limit the effect of heterogeneity in point density, the point clouds are voxelized to a 0.1 m resolution within the function competition_pc(). Depending on the selected method ("cone" or "cylinder"), the voxels that fall into the search cone or cylinder (radius/cone height adjustable) are counted as a competing neighbourhood. For these methods, the choice of the cone opening point or the cylinder diameter has a high impact on the calculated outcomes and should therefore be chosen responsibly. By default, the position of the cone opening point or the centre of the cylinder is the metroid of the CPA. The stem base point can also be selected, but then the search cone may be partially or completely outside the tree crown if the stem of the target tree is inclined.

**Figure 2:**
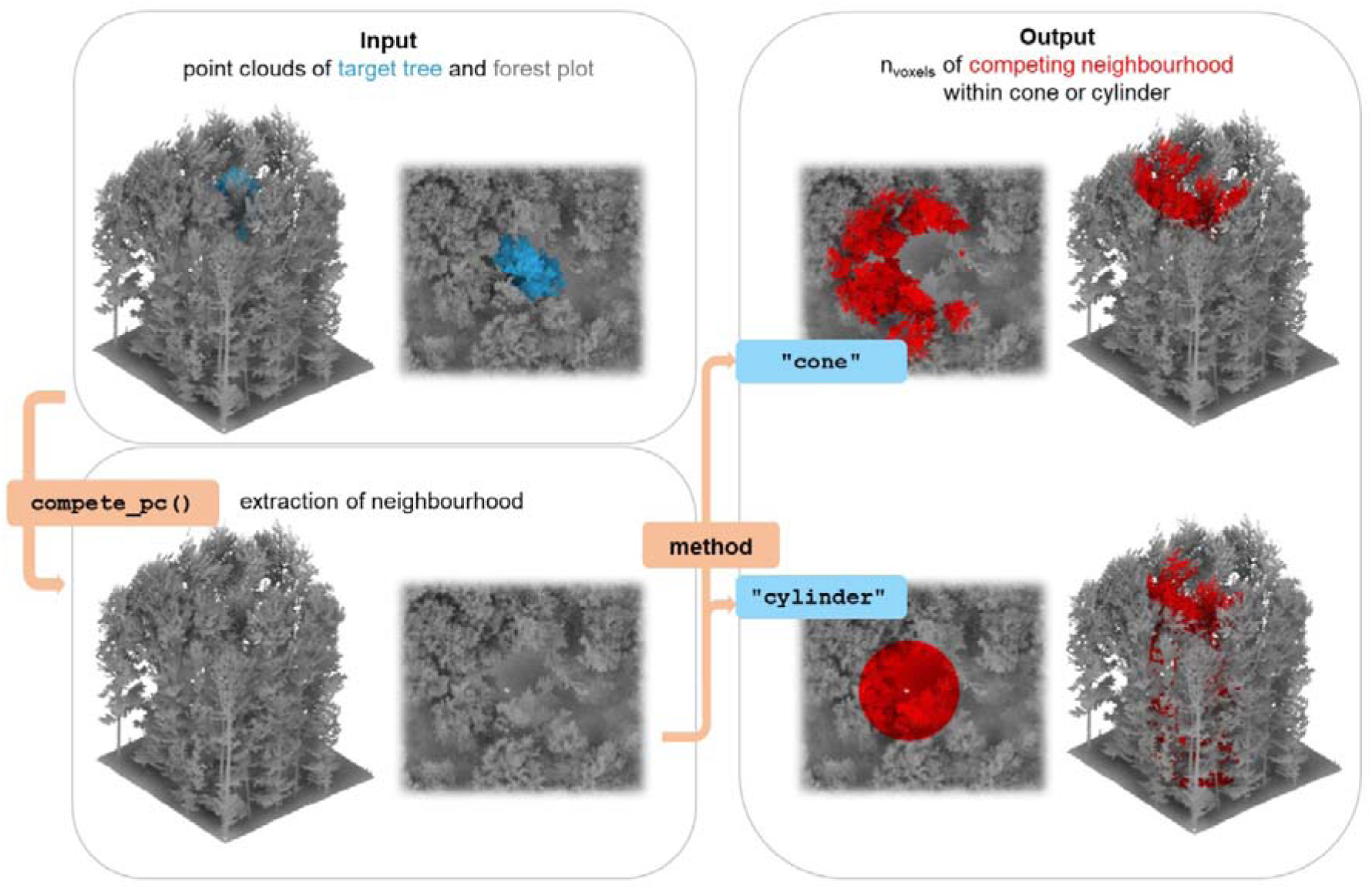
Functionality of compete_pc() with filtering of the competing neighbourhood (left) and the "cone" or "cylinder" method (right).

## 3. Results and discussion

### 3.1 Competition quantified from MLS point clouds

The 308 MLS scans of target trees were used to quantify the CIs with the cone method and cylinder method. For the cone method, we decided to use a cone with 60° opening angle spanned in 50% of each target tree’s height, as the commonly-used 60% threshold (Seidel et al., 2015) would have resulted in a large number of trees with voxel counts of zero (no apparent competition). For the cylinder method, we used a 4.5 m radius, which is the average crown diameter of our trees. In addition, we used the MLS scans to compute the CI_Hegyi_ and CI_Braathe_, both with search radius of 13.5 m assumed to reflect 3× average crown radii. A comparison of the CIs with different settings is shown in Figure 3. Due to the differences in calculation and input variables, the range of the CI values differs for each index. Besides the method, the chosen search radius plays an important role, as it defines how many trees are considered as competitors. The sensitivity to this parameter can be observed in Figure 3A, where we plot the range of the distance-dependent CIs for the different search radii 4.5 m, 9 m and 13.5 m. With increasing search radius, the CI also increases. In addition, the cone height and cylinder radius within the point cloud-based approaches have a clear influence.

**Figure 3:**
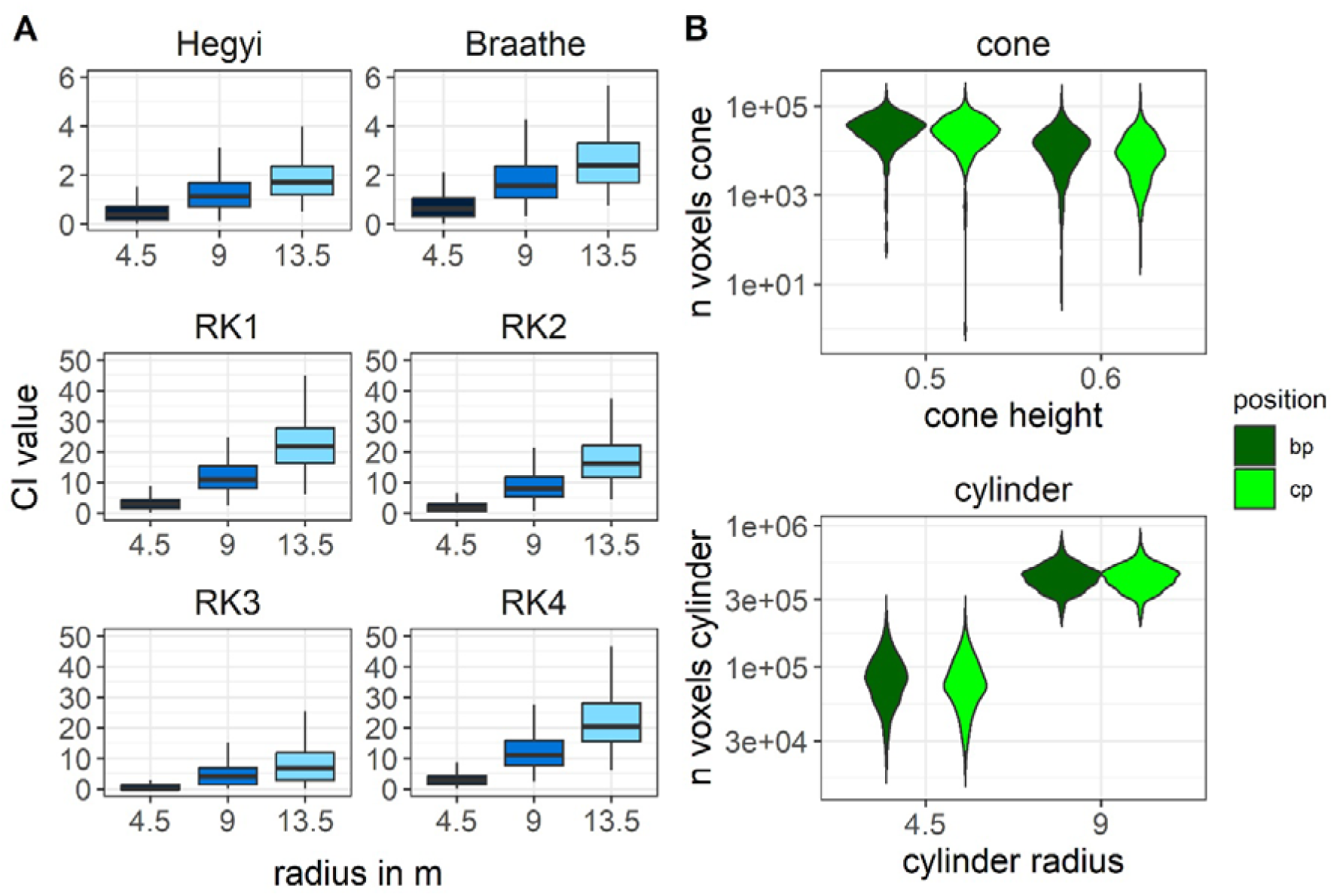
**A.** Distribution of the distance-dependent competition indices (CIs) derived from mobile laser scanning (MLS) and three different search radii (1×, 2×, 3× average crown radius of 4.5 m) for the 308 target trees. **B.** Comparison of CI values for cone and cylinder method with different cone heights (50 and 60 % of target tree’s height) and different cylinder radii (1× and 2× crown radius) and the tree position methods base position (bp) and crown position (cp)

The correlation between structural parameters, as height (H, m), diameter at breast height (DBH, m), crown projection area (CPA, m^2^), box-dimension (D_b_) and the CIs is shown in Figure 4. In general, we observed a negative linear correlation between the structural parameters and the CIs, indicating that higher competition leads to smaller crowns and lower D_b_. The strongest linear correlation with CPA was found for the distance-dependent indices CI_Hegyi_ and CI_Braathe_ (r = -0.55 and -0.38, respectively), whereas D_b_ showed the strongest correlation with CI_Braathe_ (r = -0.43) and CI_Hegyi_ (r = -0.41). The distance-dependent indices were strongly correlated with each other (r = 0.84) due to the similar computation. The CIs derived from point clouds were not significantly correlated to each other, but CI_cylinder_ was correlated a bit more strongly with the distance-dependent indices than CI_cone_. Within the *TreeCompR* package, more distance-dependent CIs are available (CI_RK1_ to CI_RK4_) but not presented here, as they are rarely used in the literature. In general, the point cloud-based CIs were more weakly correlated to structural parameters than the traditional approaches.

**Figure 4:**
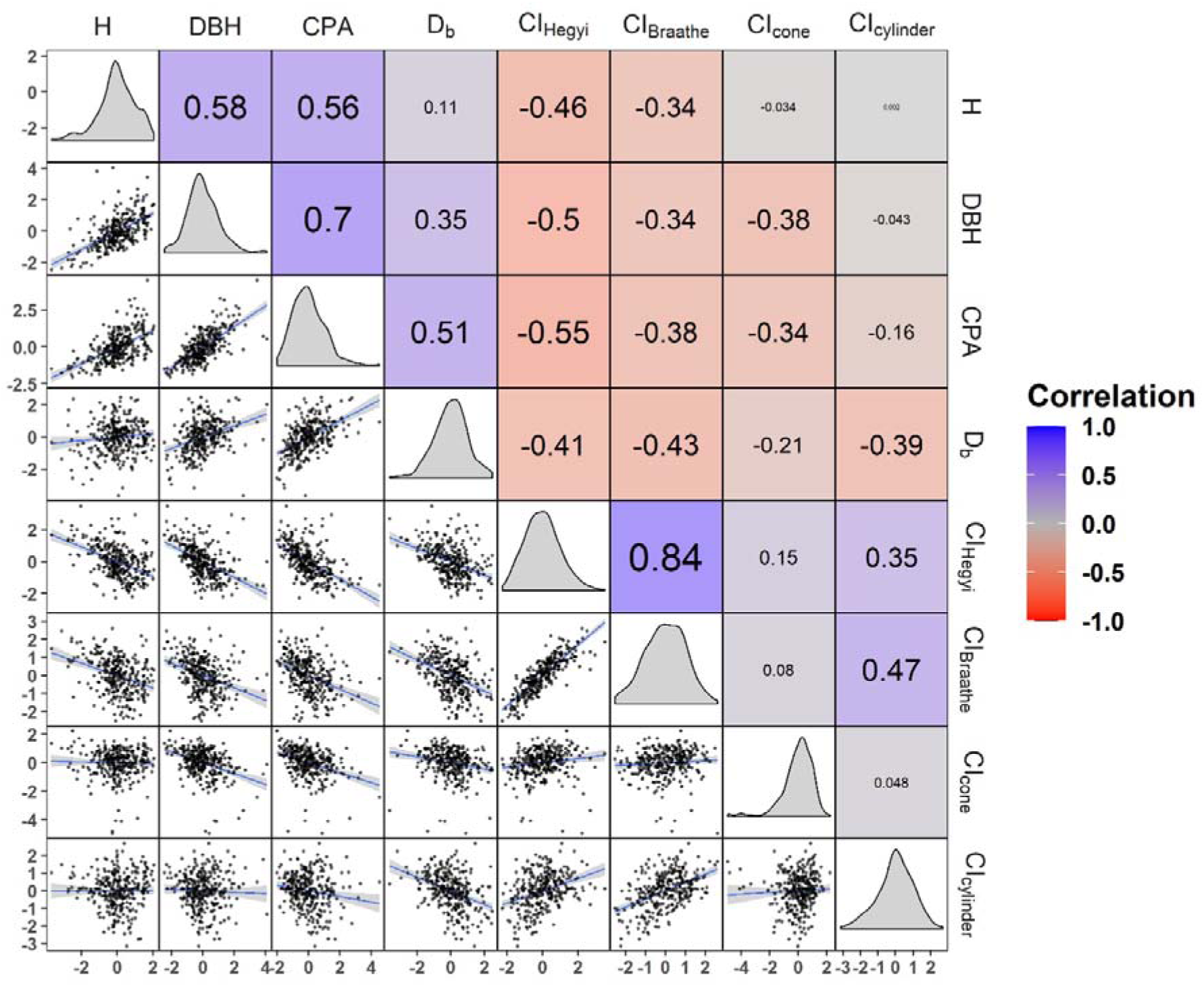
Correlation plot of target trees’ heights (H, m), diameter at breast height (DBH, m), crown projection area (CPA, m^2^), box dimension (D_b_) and the four competition indices (CIs) quantified using the *TreeCompR* package. CIs are log-transformed, all variables centred and scaled by standard deviation. Shown are the Pearson correlation coefficients (upper triangle, shaded by correlation strength), density plots of the variables (diagonal) and scatter plots overlaid with a linear model fit (blue line) with its 95 % confidence intervals (grey ribbon). Figure created using *corrmorant* v0.0.0.9007 (Link, 2020).

### 3.2 Effect of data source

We also tested ALS data to quantify the height-distance dependent CIs. CI_Braathe_ had clearly higher values in our example for ALS compared to MLS data (cf. Figure 5A). This is most likely due the difference in the identified number of neighbours within a 13.5 m radius around our target trees for the two data sources (cf. Figure 5B), with significantly higher counts for the segmented ALS data, particularly in a subset of forest sites. This most likely is due to oversegmentation artefacts and highlights that ALS or MLS data or segmentation results should be compared to ground-based information before use whenever possible to avoid biased results as higher numbers of neighbours lead to higher CI values. Finding an appropriate algorithm for segmentation depends on the type of data collection, the point density and potentially even the tree species (Lu et al., 2014). For this reason, we decided against implementing any of these methods within this package. Besides errors in the number of trees within a plot, an overestimation of DBH can occur in LiDAR-based data if branches are present at breast height, and the derived data should always be checked for outliers driven by extreme DBH values.

**Figure 5:**
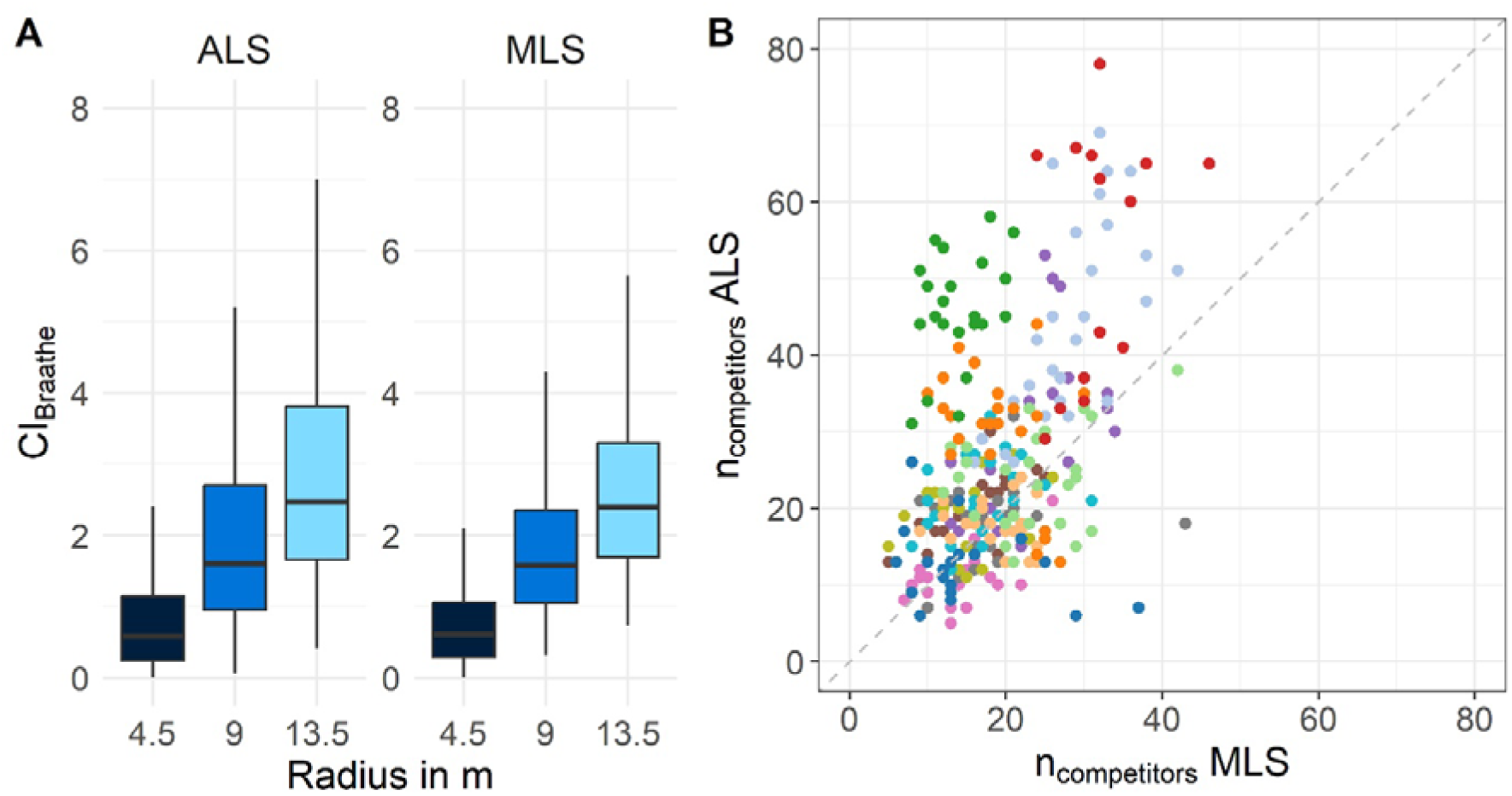
**A.** Distribution of height-distance dependent CI_Braathe_ derived from airborne (ALS) and mobile laser scanning (MLS) and three different search radii (1×, 2×, 3× average crown radius of 4.5 m) for the 308 target trees. **B.** Comparison of number of competitors within search radius of 13.5 m around the target trees derived from ALS vs. MLS data. Different colours indicate different forest sites.

Measurement errors can also occur in the field, as tree heights can be difficult or even impossible to obtain by ground-based measurements in dense forest plots. In addition, there can also be inaccuracies in measuring the distance to other trees or a tree’s position if there is no adequate GPS signal. These sources of uncertainty should be considered before choosing the method and kept in mind while evaluating the results.

For all data sources, it is important to note that differences in the number of trees may also be due to differences in the time of data collection. In our case, the ALS data was collected 8-10 years before the MLS scans and depending on the research question, some datasets may be too old as trees may have been removed and parameters may have changed.

### 3.3 Decision guidance

Most published studies that quantify tree competition based on competition indices rely on a single index, in most cases a distance-dependent one such as the Hegyi index (Versace et al., 2019; Seidel et al., 2019; Saarinen et al., 2021).We recommend that users should decide on an appropriate CI a priori based on data availability, research question and/or efficiency in data collection, as none of the available indices is universally applicable.

If comprehensive inventory data is available for the study area, it makes sense to use one of the distance-dependent indices. Similarly, close-range high-resolution 3D point clouds are recommended if a laser scanner is available and if the scans can also be used for other purposes. When deciding between distance-dependent and point cloud-based CIs, it is important to note that the manual segmentation of the target trees is much more accurate than automated segmentation. Due to the high effort required for manual segmentation, point cloud-based methods are of more interest if the individual tree point clouds can be used for additional study purposes beyond quantifying competition (e.g., for structural complexity metrics such as D_b_, or for crown parameters). In the future, automatic segmentation of target trees with the help of deep learning algorithms (cf. Allen et al., 2022) seems to be a promising route to make the processing of laser scanning data more time-efficient and less labour-intensive.

To minimize sources of error in ALS data and simultaneously reduce fieldwork time, a combination of the approaches can be useful. If accurate coordinates of the trees in the plot are available, tree heights can be extracted from a canopy height model (CHM) derived from ALS data, while the number of trees within the plot can be more accurately determined from field observations. This should improve the accuracy of measured tree heights compared to ground-based measurements, at least in denser stands, but requires a sufficient GPS signal in the forest. It is not possible to measure DBH directly using ALS data, but CI_Braathe_ can be easily derived based on the tree heights and positions.

The same index and search radius should be used if the competitive situation is to be compared between studies. In the context of the distance-dependent CIs shown, it is important to note that in these indices, no distinction is made between intra- or inter-specific competition (though they can be decomposed into species-specific components; cf. Hajek et al., 2022). In contrast, the two point-cloud-based approaches (CI_cone_, CI_cylinder_) are based on the actual shape of trees, which directly reflects inter- and intraspecific variations of tree structures (cf. Metz et al., 2013).

### 3.4 Outlook

In the context of global-change-type drought stress that meanwhile affects all forested biomes of the world (Hammond et al., 2022), understanding the competitive situation within stands is crucial. In Central Europe, there is an urgent need to understand the small-scale variability in vitality in beech, which is directly affected by the competitive situation. This species is known to be shade-tolerant and sensitive to light intensity especially during periods of drought and high temperature (Leuschner, 2020). Increased competition from direct neighbours should therefore not be seen as one-sided or linear. Indeed, Ma et al. (2023) showed that more exposed trees (less competitive pressure) were more affected by drought-induced tree mortality than those shaded by larger trees, indicating that competition during drought periods may have to be evaluated differently than during normal climatic conditions. Our package lends itself to capture these sorts of patterns by calculating competition indices species-wise to model directed species interactions (cf. Hajek et al., 2022). With the availability of large-scale ALS data and automated segmentation methods, our package further enables mapping competition intensity on the ecosystem scale.

## 4. Conclusion

In summary, the *TreeCompR* package provides a comprehensive solution for assessing tree competition in forest ecosystems and allows researchers to browse through different indices and gain insight how they are used in the literature. We provide ample guidance on how to choose an appropriate index, a decision that should be informed by data availability, time, costs, domain knowledge about the species and ecosystem and, most importantly, the research question of the study. *TreeCompR* provides a range of commonly used CIs and allows the use of diverse data sources as input. The computed output of the package is geared towards easy integration in existing data analysis workflows and facilitates downstream use of the estimated CIs in statistical models. We hope that by providing an interface to calculate a broad range of metrics in a common framework, our package contributes to a more standardized reporting and an increased comparability of competition indices in published literature.

## Author Contributions

B.S. and C.Z. received the funding and conceived the original research idea. J.S.R., K.K. and A.Ž. collected the data. J.S.R., K.K. and D.S. pre-processed the data. J.S.R. and R.M.L. wrote the R package. J.S.R. analysed the data and led the writing of the manuscript together with R.M.L.. All authors contributed critically to the drafts and gave final approval for publication.

## Acknowledgements

B.S. and C.Z. gratefully acknowledge funding by the German Research Foundation (SCHU 2935/2-1, ZA 755/2-1).

## Data Availability

Our R package is available for download on Github under the following link: https://github.com/juliarieder/TreeCompR.

